# Automatic recognition of complementary strands: Lessons regarding machine learning abilities in RNA folding

**DOI:** 10.1101/2023.04.20.537615

**Authors:** Simon Chasles, François Major

## Abstract

Prediction of RNA secondary structure from single sequences still needs substantial improvements. The application of machine learning (ML) to this problem has become increasingly popular. However, ML algorithms are prone to overfitting, limiting the ability to learn more about the inherent mechanisms governing RNA folding. It is natural to use high-capacity models when solving such a difficult task, but poor generalization is expected when too few examples are available. Here, we report the relation between capacity and performance on a fundamental related problem: determining whether two sequences are fully complementary. Our analysis focused on the impact of model architecture and capacity as well as dataset size and nature on classification accuracy. We observed that low-capacity models are better suited for learning with mislabelled training examples, while large capacities improve the ability to generalize to structurally dissimilar data. It turns out that neural networks struggle to grasp the fundamental concept of base complementarity, especially in lengthwise extrapolation context. Given a more complex task like RNA folding, it comes as no surprise that the scarcity of usable examples hurdles the applicability of machine learning techniques to this field.

## Introduction

Identifying potential structural candidates for a single RNA sequence is a computationally demanding task. The Zuker-style dynamic programming approach to fold an RNA sequence of length *L* without pseudoknots requires time complexity in 𝒪(*L*^3^) [Zuker and Stiegler(1981)] [Hofacker et al.(1994)Hofacker, Fontana, Stadler, Bonhoeffer, Tacker, Schuster et al.]. Algorithms that take into account pseudoknots are even more complex and have been reported to require significantly more computational power ranging from 𝒪(*L*^4^) to 𝒪(*L*^6^) [Rivas and Eddy(1999)] [Condon et al.(2004)Condon, Davy, Rastegari, Zhao, and Tarrant], or higher [Marchand et al.(2022)Marchand, Will, Berkemer, Bulteau, and Ponty].

Machine learning (ML) algorithms offer an alternative to traditional methods for identifying RNA structural candidates. In particular, neural networks can compute structures in an end-to-end fashion, allowing for quick inference in a single feedforward pass, especially when running on `GPU` [Chen et al.(2020)Chen, Li, Umarov, Gao, and Song] [Fu et al.(2022)Fu, Cao, Wu, Peng, Nie, and Xie]. However, the training phase can take several days, which can complicate software updates [Shen et al.(2022)Shen, Hu, Peng, Chen, Xiong, Hong et al.]. Regardless of the approach, predicting RNA structure requires significant computational resources due to the inherent complexity of RNA folding. As prediction methods must be at least as complex as the problems they aim to solve, it is important in the case of ML to avoid overfitting to this task. To match the inherent complexity of RNA structure prediction, ML algorithms require high capacity, meaning they should be capable of learning a wide variety of mathematical functions [Goodfellow et al.(2016)Goodfellow, Bengio, and Courville]. Actually, because statistical learning relies heavily on training data, ML algorithms require high finite-sample expressivity to learn effectively [Zhang et al.(2021)Zhang, Bengio, Hardt, Recht, and Vinyals]. However, high expressivity can lead to overfitting if the neural networks perform well on the training data without being able to generalize to structurally dissimilar testing data [LeCun et al.(1989)]. Therefore, it is crucial to balance expressivity with generalization to ensure accurate predictions on unseen data.

Several ML algorithms have been developed for RNA secondary structure prediction in recent years [Chen et al.(2020)Chen, Li, Umarov, Gao, and Song] [Fu et al.(2022)Fu, Cao, Wu, Peng, Nie, and Xie] [Zakov et al.(2011)Zakov, Goldberg, Elhadad, and Ziv-Ukelson] [Singh et al.(2019)Singh, Hanson, Paliwal, and Zhou] [Wang et al.(2019)Wang, Liu, Zhong, Liu, Lu, Li et al.]. However, many of these algorithms are suspected of overfitting [Rivas et al.(2012)Rivas, Lang, and Eddy] [Sato et al.(2021)Sato, Akiyama, and Sakakibara] [Zhao et al.(2021)Zhao, Zhao, Fan, Yuan, Mao, and Yao] and have limited ability to generalize across RNA families [Szikszai et al.(2022)Szikszai, Wise, Datta, Ward, and Mathews]. Generalization is crucial for accurate RNA structure prediction since known RNA structures only represent a small fraction of the entire RNA structure space. Prediction algorithms must be able to accurately predict structures for molecules that are similar and dissimilar to known structures. Therefore, it is important to develop ML algorithms with stronger generalization properties to accurately predict RNA structures across a wider range of sequences and structures.

The aim of this study is to explore the performance and behavior of ML algorithms on a fundamental RNA-related task: determining if two RNA strands are fully complementary. Specifically, we examined the behavior of four families of neural networks with a focus on overfitting. We tackled three major challenges encountered when applying ML to RNA folding: (1) learning with mislabelled training examples (mislabels), (2) generalizing to structurally dissimilar data, and (3) training with limited examples [Rivas et al.(2012)Rivas, Lang, and Eddy] [Flamm et al.(2021)Flamm, Wielach, Wolfinger, Badelt, Lorenz, and Hofacker] [Burley et al.(2022)Burley, Bhikadiya, Bi, Bittrich, Chen, Crichlow et al.] [Danaee et al.(2018)Danaee, Rouches, Wiley, Deng, Huang, and Hendrix].

Our results indicate that low-capacity models are more effective for learning with mislabels, as they have the ability to ignore them. Conversely, high-capacity models demonstrate better generalization performance in length-wise extrapolation context. On top of that, learning with few examples poses challenges for both low and high-capacity models, highlighting the importance of problem representation and architecture choice.

## Materials and methods

### Learning task

Given the RNA alphabet Σ = `{A, C, G, U}`, complementary strands are those in which each nucleotide on one strand pairs with its Watson-Crick partner on the other strand (`A` with `U` and `C` with `G`). For example, the RNA strand 5’-`AGUCAG`-3’ is complementary to 5’-`CUGACU`-3’. We defined the task of automatic recognition of complementary strands as a binary classification problem that involves comparing and determining whether pairs of RNA strands of the same length are complementary or not. The target is `True` if the strands are fully complementary and `False` otherwise.

To have a fixed-size noisy input and simulate the structure of a hairpin loop, we restricted the maximum length of an RNA strand to 8 nucleotides and inserted an apical loop of 4 random nucleotides between each pair of RNA strands. More specifically, given strands *s*, 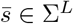 with *L ≤* 8 and *a ∈* Σ^4^, we represented the input as the sequence 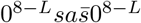, where 0 denotes zero-padding.

The target of the classification is *t* = 1 if for all *i ∈ {*1, …, *L}, s*_*i*_ is paired with its Watson-Crick complement on the other strand, that is, if 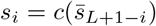, where *c* : Σ *→* Σ is the Watson-Crick complementarity function defined as *c*(`A`) = `U`, *c*`(C)` = `G`, *c*`(G)` = `C`, and *c*`(U)` = `A`. Otherwise, the target is *t* = 0.

Each nucleotide was represented by a vector of size 4, and a word embedding layer was used to update these vectors during training. Therefore, the inputs are fixed-size real-valued matrices *x ∈* ℝ^20*×*4^ and the outputs are real values *y ∈* (0, 1). The output can be interpreted as the probability of the RNA sequence being a positive example, which we also refer to as the positivity score.

The loss function used to train the models is the binary cross-entropy with adjustments made to account for varying ratios of positive and negative examples.Specifically, for a training dataset 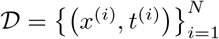 with positive example ratio 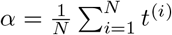, the loss value for a model prediction *y*^(*i*)^ = *f* (*x*^(*i*)^) was computed by Eq (1).

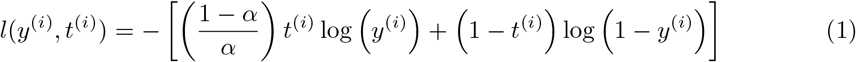

This correction ensured that the loss for positive examples was scaled up or down relative to the loss for negative examples, subject to the dataset’s positive example ratio. For instance, if there were twice as many negative as positive examples in a dataset (i.e., *α*= 1/3), the loss value for each positive example would be multiplied by a factor of 2, effectively balancing the contribution of positive and negative examples to the training loss. The parameter *α* could be controlled when creating synthetic datasets.

### Artificial data

Our data generation process aimed to create diverse training and testing datasets that would enable us to evaluate our models in various scenarios. To achieve this, we introduced structural and statistical dissimilarities between the datasets, including differences in quality, sequence length and positivity rate. Mainly, we aimed at simulating the use of all known data (a training set) to infer predictions on a portion of the unseen data of interest (a corresponding testing set).

To produce a pair of training and testing datasets, we ensured that there was no overlap between the training and testing examples by randomly partitioning the set of all 4^*L*^ sequences of length *L* into two sets. For each example, we concatenated a sequence *s ∈* Σ^*L*^ with a randomly generated apical loop *a* and a complementary sequence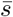. Specifically, for positive examples, we set 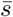 to be the exact complement of *s* (denoted by *s*^*\**^), while for negative examples, we chose 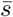randomly in Σ^*L*^ *\ {s*^*\**^*}*. We carefully controlled the ratio *α* of positive examples in the training sets and ensured that the testing sets had an equal number of positive and negative examples, thereby avoiding bias when measuring the performance on a test set.

To introduce structural dissimilarities, we varied the sequence length and the positive example ratio between the training and testing datasets. This allowed us to measure the extrapolation abilities of our models. Additionally, to control the quality of a training dataset, we introduced a proportion 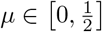 of mislabelled examples, where a positive example could have label 0 with probability *µ*, and a negative example could have label 1 with probability *µ*. However, we ensured that all examples in the testing datasets were correctly labelled. By doing so, we tested the models’ ability to ignore errors and avoid overfitting.

Overall, our data generation process produced realistic and diverse datasets, allowing us to evaluate our models under various conditions. Fully aware of the similarity between the task defined here and the prediction of binding sites of microRNAs, we are mostly interested in the abilities and behaviors of ML algorithms (and neural networks in particular) when trained and tested in various conditions, which we better control using artificial data.

### Performance measure

To evaluate the performance of our models, we measured their classification accuracy on the testing datasets using a classification threshold *θ ∈* (0, 1) to distinguish between examples of class 0 and 1. Specifically, given a model *f* (*·*) and a dataset 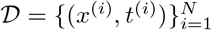, the accuracy of *f* (*·*) on 𝒟 with threshold *θ* was calculated by Eq (2).

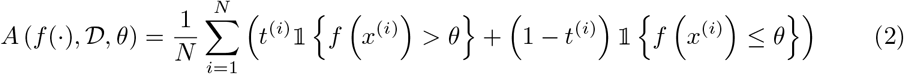

where 𝕝*{·}* is the indicator function that equals 1 if the condition inside the bracket is true, and 0 otherwise.

While threshold *θ* could be optimized through methods such as SVM [Smola and Schölkopf(2004)] or maximizing accuracy on the training dataset [Singh et al.(2019)Singh, Hanson, Paliwal, and Zhou], we set *θ* = 1*/*2 to study the behavior of our neural networks independently of any additional optimization steps. We considered the mean accuracy over 50 simulations, each of which generated datasets as described above, trained a model as described in the next section, and evaluated its performance using the aforementioned accuracy metric.

### Architectures and training

The four tested models are relatively small representatives of four types of neural networks that have been recently used to predict RNA structures [Chen et al.(2020)Chen, Li, Umarov, Gao, and Song] [Singh et al.(2019)Singh, Hanson, Paliwal, and Zhou] [Sato et al.(2021)Sato, Akiyama, and Sakakibara]. All models have only three layers, where the first layer is unique to each architecture and the last two layers have the form `linear - batchnorm - ReLU - dropout - linear-sigmoid` [Goodfellow et al.(2016)Goodfellow, Bengio, and Courville] [Ioffe and Szegedy(2015)] [Nair and Hinton(2010)] [Hinton et al.(2012)Hinton, Srivastava, Krizhevsky, Sutskever, and Salakhutdinov]. We used a dropout rate of 0.1 and a weight decay of 10^−3^ for all models, and the number of epochs was set to 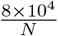 to ensure that all models were trained for the same number of iterations, regardless of the number of training examples *N*. We used the `Adam` optimizer with a learning rate of 10^−3^ and default parameters in `PyTorch` [Kingma and Ba(2014)] with a batch size of 256.

The four tested models and their capacity control are summarized in Fig 1. The multi-layer perceptron model (**MLP**) [Goodfellow et al.(2016)Goodfellow, Bengio, and Courville] has a first layer of the form `linear - batchnorm - ReLU - dropout`, and its capacity is controlled by the number *H*_1_ of hidden units in the linear module. For the multi-head self-attention model (**Att**), the first layer uses a skip connection of the form *h*_1_ = conv(*x* + positional encoding) and *h*_2_ = batchnorm(*h*_1_ + dropout(multi-head self-attention(*h*_1_))), where the multi-head attention and positional encoding are described by Vaswani and co-workers [Vaswani et al.(2017)Vaswani, Shazeer, Parmar, Uszkoreit, Jones, Gomez et al.]. The capacity of this layer is controlled by the number *H*_2_ of heads in the multi-head attention, so the 1D-convolution uses a kernel of size 1 to project the 4-dimensional embedding into a *H*_2_-dimensional input for the attention module. For the long-short term memory model (**LSTM**) [Hochreiter and Schmidhuber(1997)] [Sak et al.(2014)Sak, Senior, and Beaufays], the first layer is a one-layer bidirectional LSTM without dropout, and its capacity is controlled by the number *H*_3_ of hidden units in the LSTM module. Finally, for the convolutional neural network (**CNN**) [LeCun et al.(1989)] [Dumoulin and Visin(2016)], the first layer has the form `outer concatenation - 3×3conv - batchnorm - ReLU - dropout` with a stride and padding of size 1. The outer concatenation operation takes the 20 *×* 4 input matrix and returns a 20 *×* 20 *×* 8 tensor, where the 8-dimensional vector at position *i, j* is the concatenation of the 4-dimensional encodings at positions *i* and *j* in the initial matrix.

**Fig 1.**
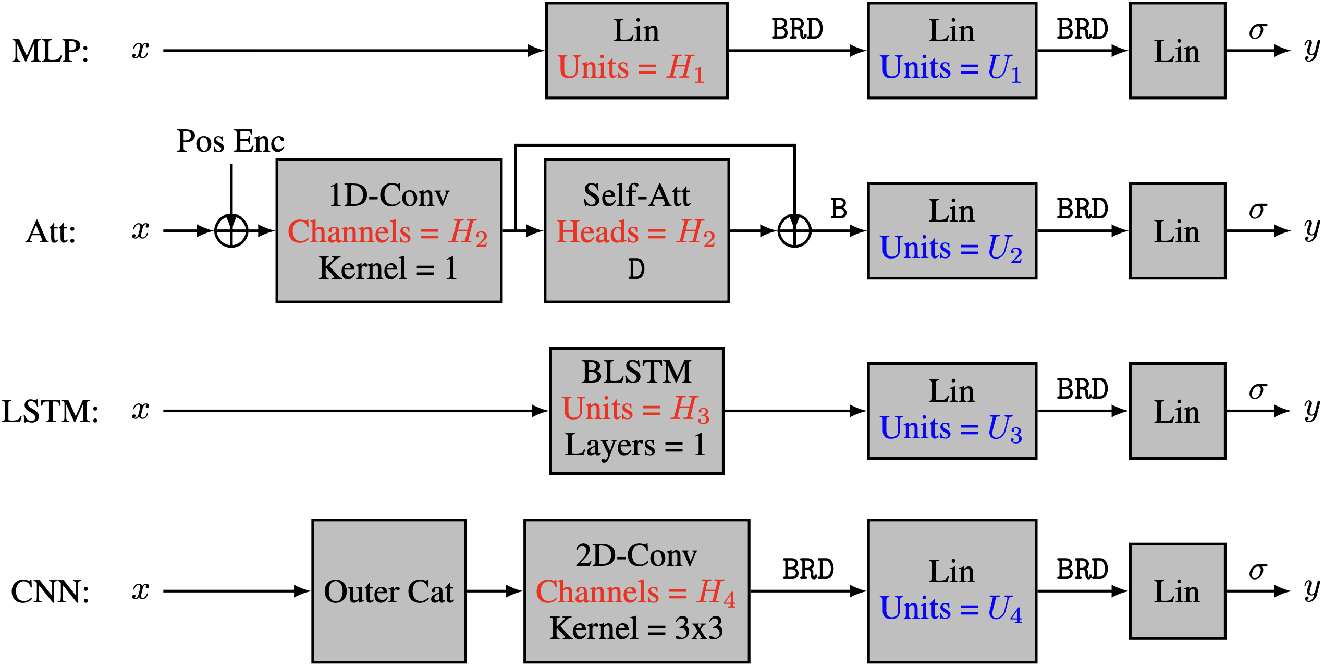
The four neural network architectures. The 4 tested neural network architectures take nucleotide encodings as input and output the positivity score. All models have three layers, with the first layer being a characteristic layer, and the last two layers having the form Lin-`B-R-D`-Lin-σ, where Lin refers to a linear layer, Conv refers to a convolutional layer, Self-Att refers to a multi-head self-attention layer, BLSTM refers to a bidirectional long-short term memory layer, Pos Enc refers to positional encodings, and letters σ, `B`, `R` and `D` refer respectively to sigmoid activation, batch normalization, ReLU activation and dropout regularization. The capacity of the models is controlled by hyperparameters *H*_*i*_ and *U*_*i*_.

The capacity of the CNN is controlled by the number *H*_4_ of feature maps produced by the convolution module.

To further manipulate the capacity of the models, we also varied the number *U*_*i*_ of hidden units in the second layer of our four architectures. Our main interest lies in the number *C* of parameters in each model, and more specifically in log_10_ *C*. The actual number of heads, hidden units, and feature maps for each value of *C* is provided in Table 1. As discussed earlier, we trained 50 instances of each model on 50 different training datasets and reported the corresponding mean test accuracy. Our goal was to investigate how the test accuracy is affected by *N* and *C* for different values of *L, µ* and α.

**Table 1.**
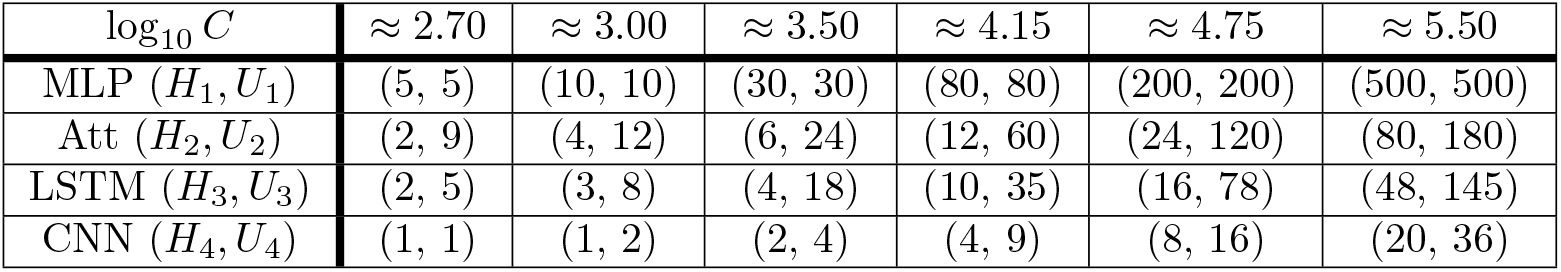
Hyper-parameters used to control the capacity of each model.

*H*_*i*_ refers to the capacity of the characteristic layer and *U*_*i*_ indicates the number of units in the second to last linear layer. The quantity log_10_ *C* gives an order of magnitude for the number *C* of trainable parameters.

## Results and discussion

### Learning with mislabels

The concept of mislabelling introduced in this section can be seen as a form of double standard. Our goal was to complicate the learning process deliberately by labelling some positive examples as class 0 and some negative examples as class 1. Thus, the term mislabelling is used because a ground truth classification is defined based on sequence complementarity. However, mislabelling on its own is not necessarily a problem for the learning process. A model *f* (*·*) that generalizes well on a completely mislabelled dataset could perform equally well on a correctly labelled dataset sampled from the same distribution by simply outputting 1 *− f* (*·*). Therefore, the learning process is complicated when a proportion *µ ∈* (0, 1) of the training dataset is mislabelled, creating a double standard as different pairs of complementary strands are labelled differently.

Moreover, since the labels can be interchanged without loss of generality, we focused on mislabelling probabilities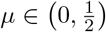. The special case where 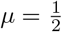behaves as if the labels were randomly assigned [Zhang et al.(2021)Zhang, Bengio, Hardt, Recht, and Vinyals]. In this study, we used a mislabelling probability of *µ* = 0.2, which we find to be a good representative of the key findings presented in this section.

In comparison to the actual RNA structure prediction task, learning with mislabels can be seen as analogous to learning on a dataset that contains structures with non-canonical base pairs as well as structures within which non-canonical base pairs are ignored, even though such pairs exist and can be identified. Similarly, the same concept applies when training on a dataset aimed at predicting pseudoknots and triplets when these kinds of interactions are only reported for some structures but not for others. Without defining the structural elements that need to be predicted, situations of double standard can arise when certain types of base pairs can be labelled as present (positive) as well as absent (negative) in a single dataset.

We were interested in investigating how the number of parameters *C* and the number of training examples *N* affect the test accuracy of the models for the automatic recognition of complementary strands. In particular, we focused on fixed sequence length *L* = 8 and mislabelling probability *µ* = 0.2. The results for the MLP and Att models are presented in Fig 2, where heatmaps depict the performance of the models for different values of *C* and *N*. Shades of red depict train accuracies; blue testing. See S1 Fig for the equivalent LSTM and CNN results. As expected, the accuracy increases with *N*, and overfitting behavior is observable for log_10_ *C ≥* 4.15.

**Fig 2.**
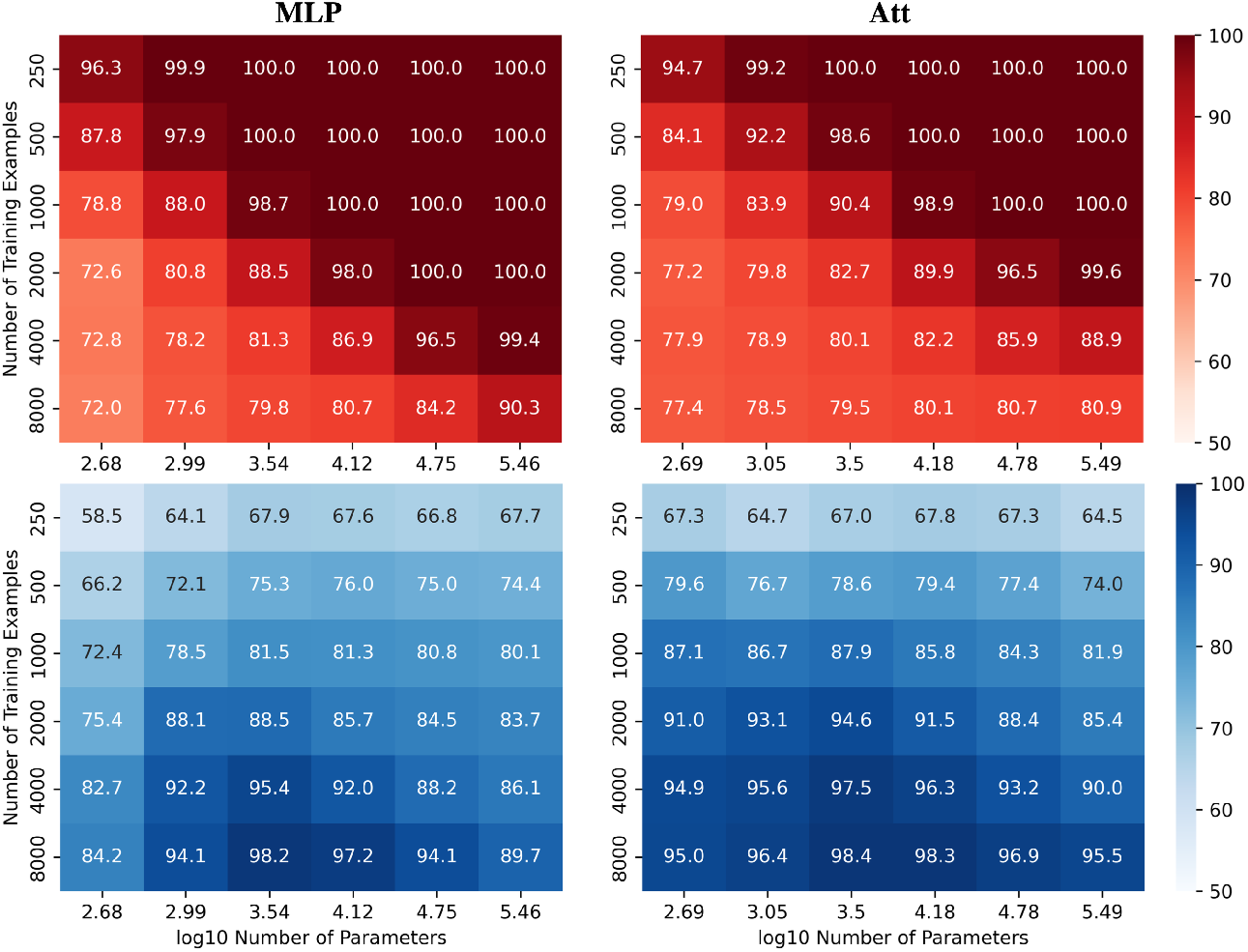
Performance of MLP and Att models when learning with mislabels. Train (red) and test (blue) mean accuracies over 50 simulations reported for MLP and Att models. Sequence length and mislabelling probability are respectively fixed to *L* = 8 and *µ* = 0.2.

Remarkably, these results demonstrate the models’ ability to handle mislabels, as they achieve a test accuracy of over 80% even when 20% of the training set is mislabelled. Moreover, the models can achieve near-perfect test accuracies as long as they are trained on a sufficiently large dataset. This finding suggests that models with relatively low capacity can still learn effectively when trained on low-quality high-quantity datasets.

Indeed, it appears that low-capacity models (log_10_ *C ≈* 3.5) are more likely to achieve test accuracies above 1 *− µ* than high-capacity models (log_10_ *C ≈* 5.5). This is likely due to the fact that mislabels introduce irregularities into the sample space that are difficult for low-capacity models to account for. Low-capacity models tend to compute smoother functions than high-capacity models. They are thus less capable of capturing the intricate patterns that arise from mislabelling, especially in large training datasets. They have however enough capacity to estimate the function of interest, yielding test accuracies near 100% and train accuracies around 1 *− µ*.

On the other hand, high-capacity models can account for the irregularities introduced by the mislabels for larger training datasets, delaying the cross-over point of train and test accuracies to larger *N*. They thus require a broader view of the sample space to attain high test accuracies, coupled with sufficient regularization to prevent overfitting. The graphs in Fig 3 illustrate these behaviors. As in Fig 2, shades of red and blue depict train and test accuracies respectively.

**Fig 3.**
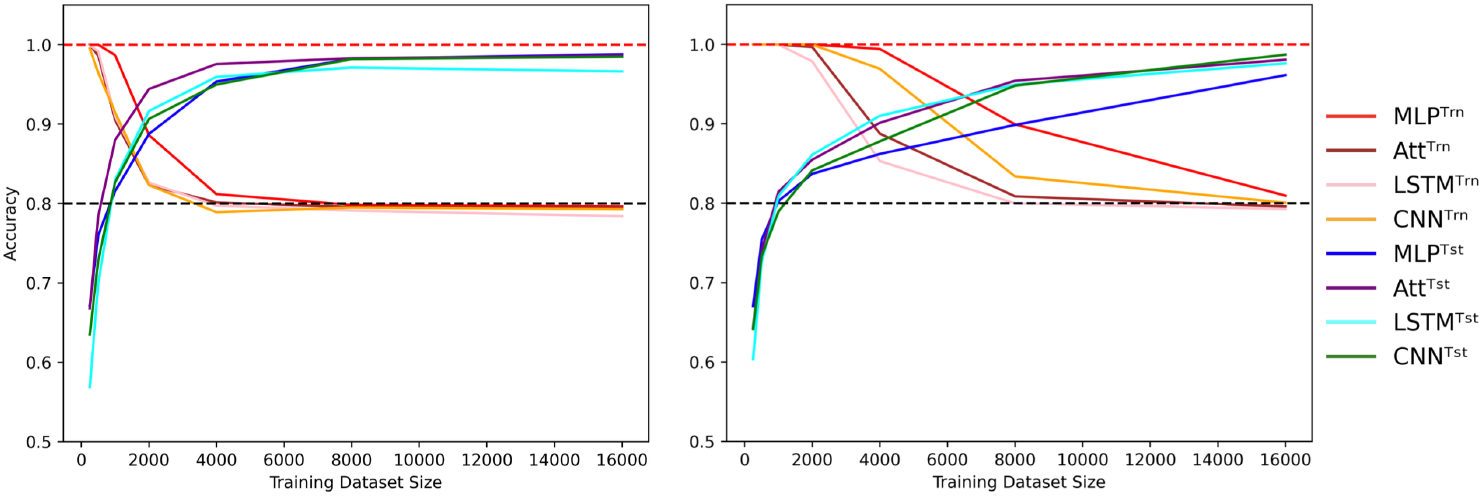
Cross-over behavior when learning with mislabels. Influence of the training dataset size over train (Trn) and test (Tst) accuracies for low-capacity models (left) and high-capacity models (right). Sequence length is fixed to *L* = 8 and mislabelling probability is fixed to *µ* = 0.2, with low capacity meaning log_10_ *C ≈* 3.5 and high capacity meaning log_10_ *C ≈* 5.5. Dotted lines indicate the 100% and 80% accuracy marks since 20% of the training examples are mislabelled.

However, it is important to note that these reported accuracies may be too optimistic because the models were trained and tested on sequences of the same length with the same ratios of positive and negative examples. Although no training sequence was repeated in the testing set, this setup only evaluates a model’s ability to generalize within structurally similar data, but not to extrapolate to structurally dissimilar data. While this section highlights the effectiveness of low-capacity models in learning with mislabels, the following section presents a contrast as high-capacity models appear to be more suitable for generalizing to structurally dissimilar data.

### Generalizing to structurally dissimilar data

In the context of the automatic recognition of complementary strands, the ability to generalize to structurally dissimilar data refers to the ability of a model to make accurate predictions over a subspace of the sample space that it has not or poorly seen during training. This means that models could be trained and tested on datasets containing different sequence lengths and positivity rates to evaluate their understanding of sequence complementarity. For example, a model can be trained on sequences of length 5 or 6 and then tested on sequences of length 8, or trained on datasets with few or a lot of positive examples and then tested on balanced datasets. This approach allows us to investigate how well a model can generalize and extrapolate its understanding of the problem. In light of current discussions regarding extrapolation in ML conditions [Berrada et al.(2020)Berrada, Zisserman, and Kumar] [Balestriero et al.(2021)Balestriero, Pesenti, and LeCun], the concept of extrapolation is used here to convey how sequences of length 8 can’t belong to the convex hull of a set of sequences of length 6 with zero-padding.

In contrast, ML algorithms used for predicting actual RNA structure face different challenges. For instance, hardware limitations may restrict the maximum length of training sequences, but the model is still expected to predict the structure of longer sequences. The presence of non-canonical base pairs, pseudoknots, and base triples can increase the number of base pairs per nucleotide, making it difficult for models trained on sequences with a lower base pair density to generalize to more sophisticated structures. Moreover, different RNA families may have varying frequencies of certain structural motifs [Moore(1999)], further complicating the generalization of models to unseen families. Despite these challenges, the ultimate goal of predicting RNA structure for all families remains the same, regardless of their structural similarity to previously known RNA structures.

To better visualize the impact of training example count *N* and the number of model parameters *C* on test accuracies for the automatic recognition of complementary strands, we report experiments on CNN and Att models. Specifically, the models were trained on correctly labelled sequences of length 6 and then tested on the full set of 4^8^ sequences of length 8. We present heatmaps (Fig 4) that illustrate the performance of the models as the number of parameters and training examples are varied. See S2 Fig for the equivalent MLP and LSTM results. We observed that higher model capacity tends to yield better performance, regardless of the number of training examples.

**Fig 4.**
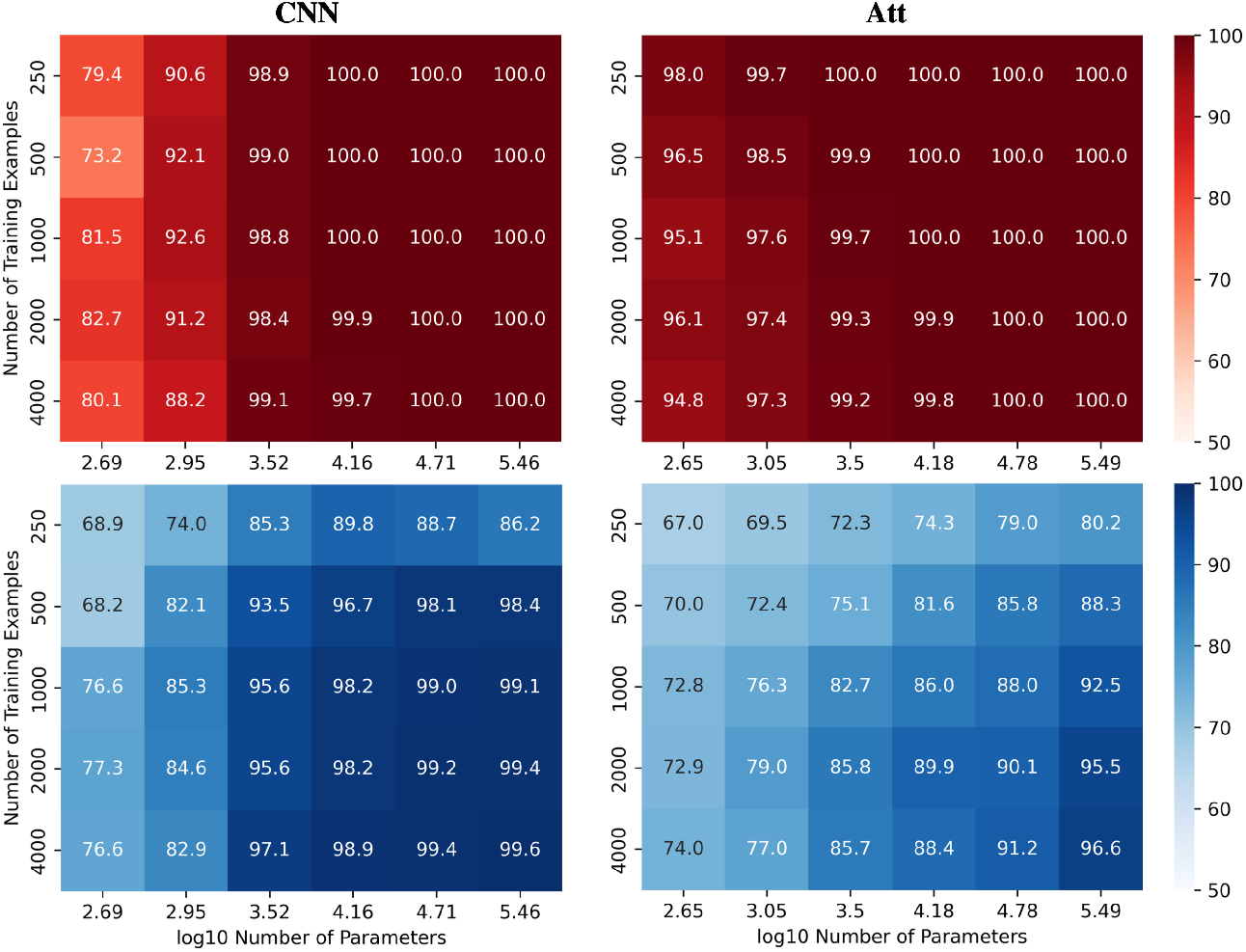
Performance of CNN and Att models in length-wise extrapolation context. Train (red) and test (blue) mean accuracies over 50 simulations reported for CNN and Att models. Models were trained on sequences of length 6 before being tested on sequences of length 8, without mislabelling in the training set.

Nevertheless, as we increase the capacity, signs of overfitting can still be observed. Notably, in the graphs presented in Fig 5, we observe signs of overfitting from the MLP model when log_10_ *C ≥* 4, for both (*L, N*) = (5, 500) on the left and (*L, N*) = (6, 2000) on the right.

**Fig 5.**
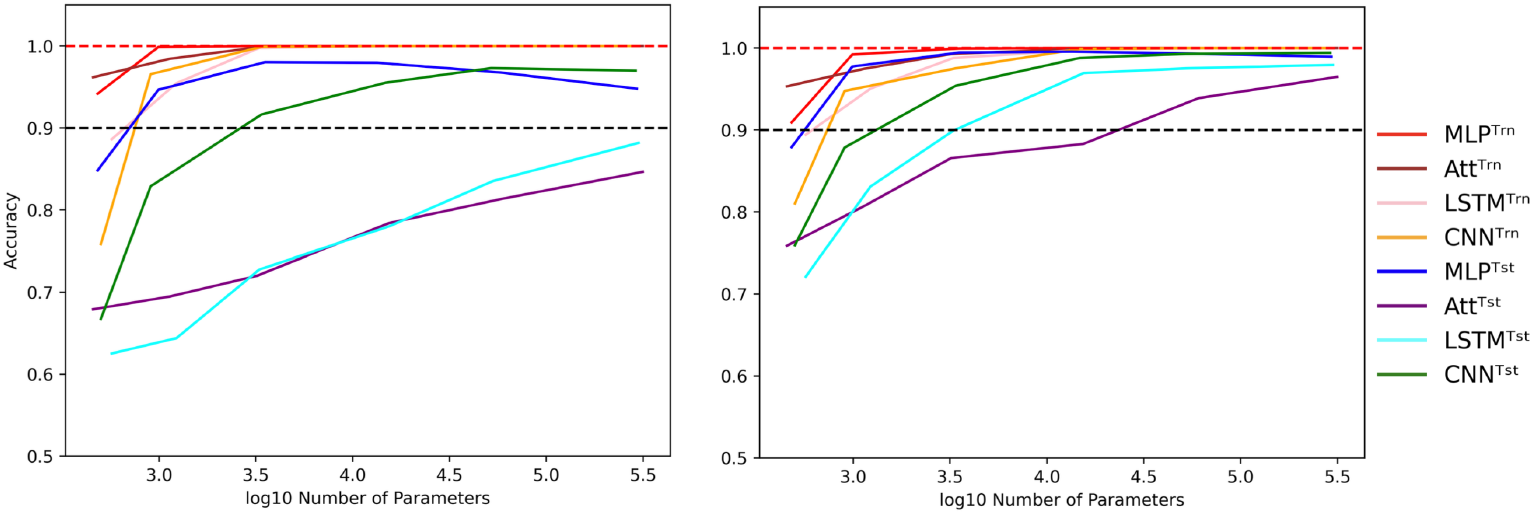
Influence of capacity in length-wise extrapolation context. Impact of the number of trainable parameters over train (Trn) and test (Tst) accuracies in length-wise extrapolation context. Results when training with 500 examples of length 5 is shown on the left and 2000 examples of length 6 on the right. Sequence length is set to 8 in the testing datasets and mislabelling probability is set to 0 in the training datasets. Dotted lines indicate the 100% and 90% accuracy marks to highlight acceptable test performance.

Interestingly, we observe that the MLP and CNN models appear to be better suited for extrapolation than the Att and LSTM models. The former can achieve test accuracies of over 90% with ease, while the latter struggle to do so. Additionally, the Att and LSTM models exhibit greater variability in their performances. For instance, the Att model of capacity log_10_ *C ≈* 4.15 with (*L, N*) = (6, 2000) has a mean test accuracy of 84.8% over 50 simulations. However, its best accuracy is over 99% and its worst accuracy is around 50%. In Fig 5, the error bars have been omitted to better highlight the performance of the four models.

Despite the impressive statistical performance of the MLP model, it fails to capture the nuances that make two RNA sequences complementary. Specifically, when trained on 500 sequences of length 5, the MLP model tends to mistake a negative example with mismatches in the upper three base pairs for a positive example. The MLP model with log_10_ *C ≈* 3.5 achieves a statistical accuracy of 98.1%, yet it incorrectly classifies the sequence ACGUACGUGAAAACGUAGCA as a positive example with a mean positivity score of 0.9810 (Fig 6). Interestingly, all other models exhibit a similar trend, as they assign a comparable (albeit slightly lower) mean positivity score to this negative sequence as they do to its corresponding positive sequence.

**Fig 6.**
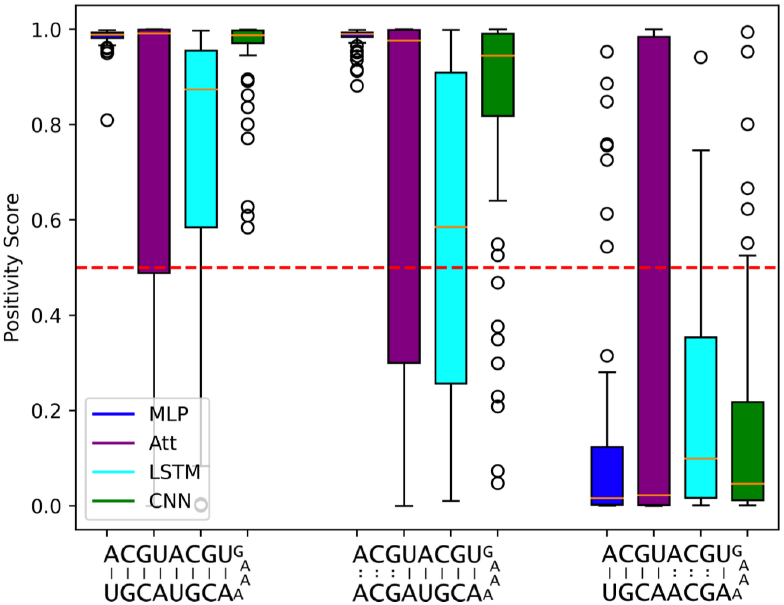
Limits to length-wise extrapolation with zero-padding and fixed-size inputs. Distribution of model outputs (positivity score) on specific sequences of length 8 when trained on 500 sequences of length 5. The first sequence is a positive example and the second and third ones are the same positive example where 3 mismatches have been introduced respectively at base pair positions 6 to 8 and 2 to 4. We use the best tested capacity for each model so log_10_ *C ≈* 3.5 for the MLP and log_10_ *C ≈* 5.5 for all other models. The classification threshold is represented by the red dotted line.

Furthermore, the negative example with mismatches in the lower five base pair positions is correctly classified most of the time by all models. This suggests that the use of zero-padding to have fixed-size intputs limits the length-wise extrapolation abilities of ML models to the nucleotide positions seen during training. The MLP model seems to be particularly affected by the zero-padding, while the LSTM tends to produce lower positivity scores for the negative sequence with mismatches in the upper three base pair positions, probably due to its sequential calculation.

It is worth noting that all results presented so far were obtained by training and testing on datasets with an equal number of positive and negative examples. However, we can manipulate the proportion *α* of positive examples in the training dataset while still testing on a balanced dataset. Despite the loss function correction that accounts for an equal number of positive and negative examples, when the concentration of positive examples is too low or too high, all four models perform poorly (Fig 7); even when classification threshold *θ* is equal to α.

**Fig 7.**
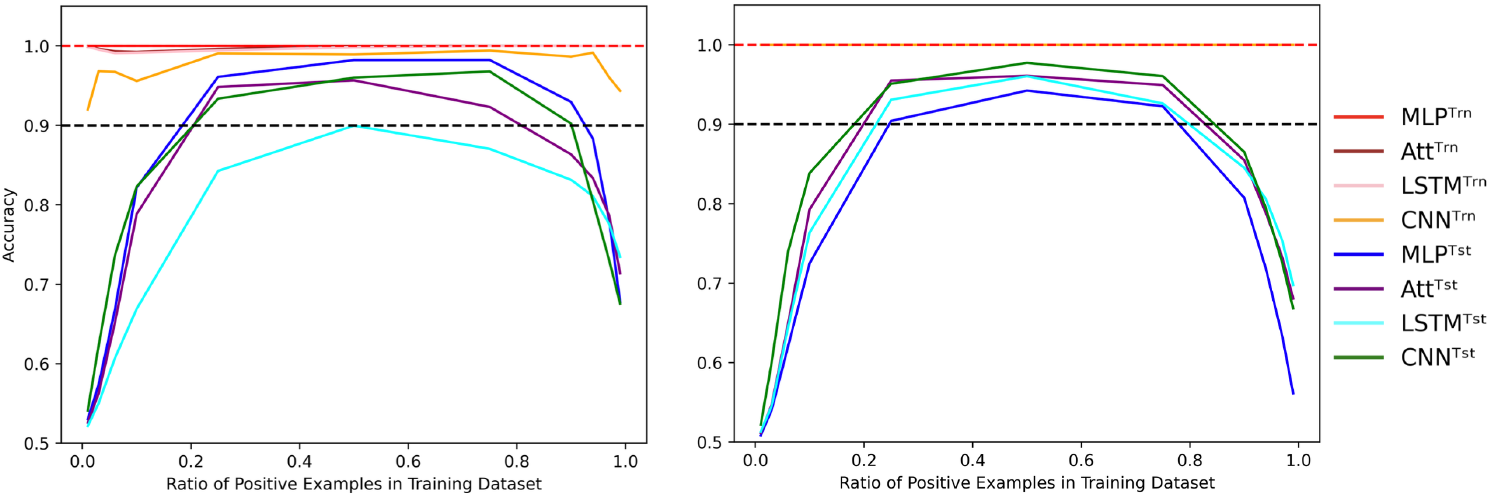
Evolution of performance in the extrapolation context of various positivity rates. Influence of the concentration of positive examples in the training set on the train (Trn) and test (Tst) accuracies for low-capacity models (left) and high-capacity models (right). Parameters were fixed to (*N, L, µ*) = (500, 6, 0). Dotted lines indicate the 100% and 90% accuracy marks to highlight acceptable test performance.

If we aim to extrapolate accurately without mislabels, high-capacity models are favorable, which means that the training datasets must be sufficiently large to support this level of expressiveness. Moreover, when our training dataset contains mislabelling, the models require a substantial number of training examples before they can disregard the errors. In the next section, we will address the challenge of learning from a small training dataset while also consolidating the insights from the previous two sections on capacity.

### Learning with few training examples

Here we focus on learning with small training datasets, but this at the same time provides us with an opportunity to explore how all our initial challenges interact with each other, both for neural networks and some classical ML methods. Thus, it is an important section that consolidates the insights gained from the previous sections.

We can note from last sections that learning with high capacity is advantageous for extrapolating without mislabels, while learning with low capacity is necessary to account for mislabels. However, when both challenges are combined, model behaviors become more complex.

To illustrate this complexity, we present heatmaps showing the behavior of the four tested architectures when trained with various quantities of sequences of length 6 with a mislabelling probability of 20% and a positivity ratio of 40% (Fig 8), the later being an arbitrary value to further challenge the extrapolation abilities of the models. The test accuracies are reported over the whole set of sequences of length 8 with as many positive and negative examples. See S3 Fig for the corresponding train accuracies.

**Fig 8.**
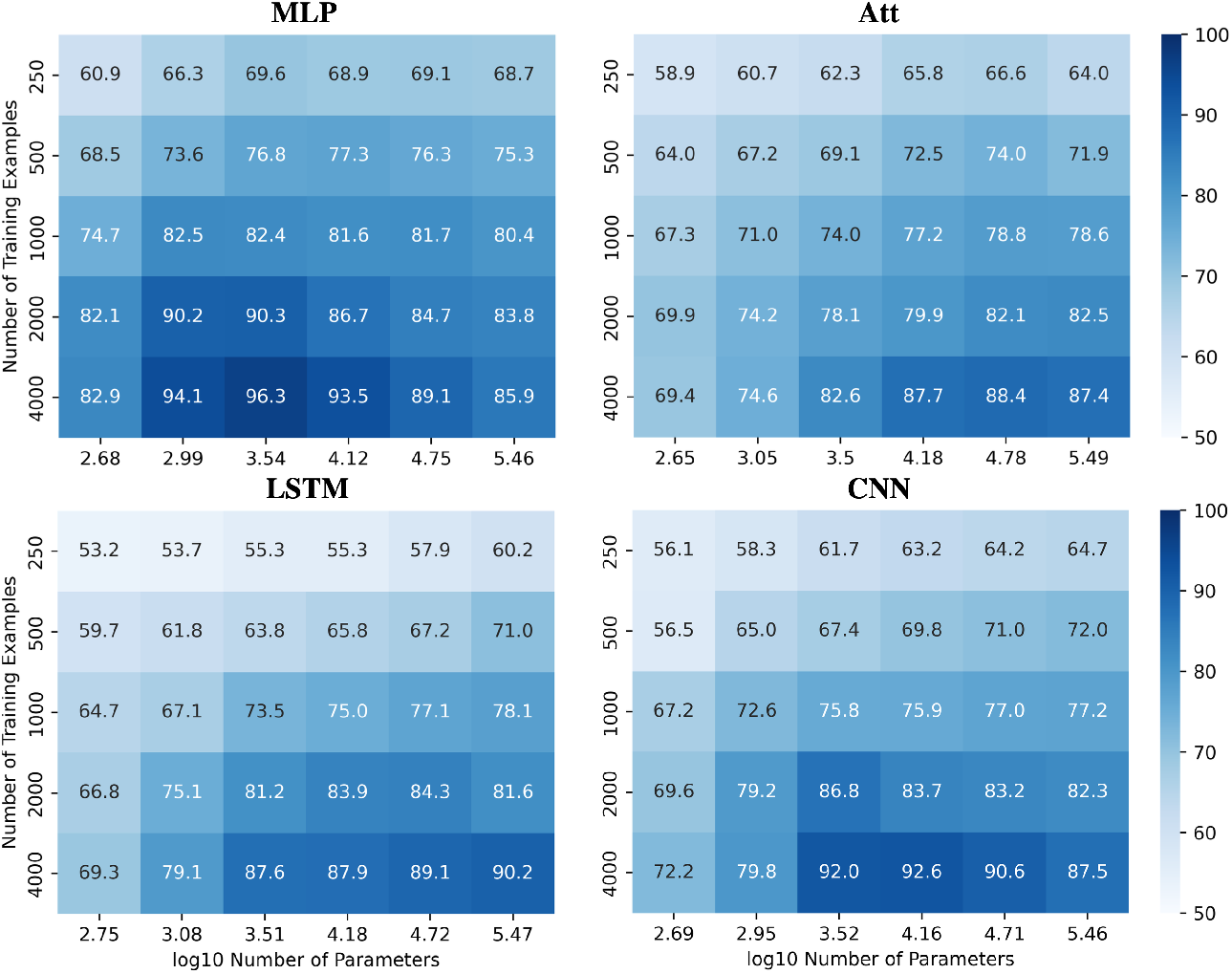
Performances for all models when learning with mislabels in extrapolation context. Test accuracies for all models when tested on sequences of length 8 after being trained on sequences of length 6 with 20% mislabelled training examples and 40% positivity rate.

These heatmaps demonstrate that the ability to ignore errors is conserved even when extrapolating, provided enough training examples are supplied. However, the required capacity to maximize generalization performance varies among the models and can be influenced by the number of training examples. For instance, the MLP and CNN require low capacity to attain their best generalization performance, while the Att and LSTM require high capacity. Additionally, the CNN’s best generalization performance requires high capacity with fewer training examples, but low capacity with more training examples.

However, achieving acceptable performances with test accuracy over 80% requires a substantial number of training examples (*N ≥* 2000) when extrapolating with mislabels. When only a few training examples are available (*N ≤* 500), test accuracies over 80% become challenging to achieve. Fig 9A shows the influence of capacity on the accuracies when trained on 500 sequences of length 6 with 20% mislabelling and 40% positivity rate, tested on the whole balanced set of correctly labelled sequences of length 8. It appears that variations in capacity have a low impact on the model’s performance, although a general slight improvement can be observed as capacity increases.

**Fig 9.**
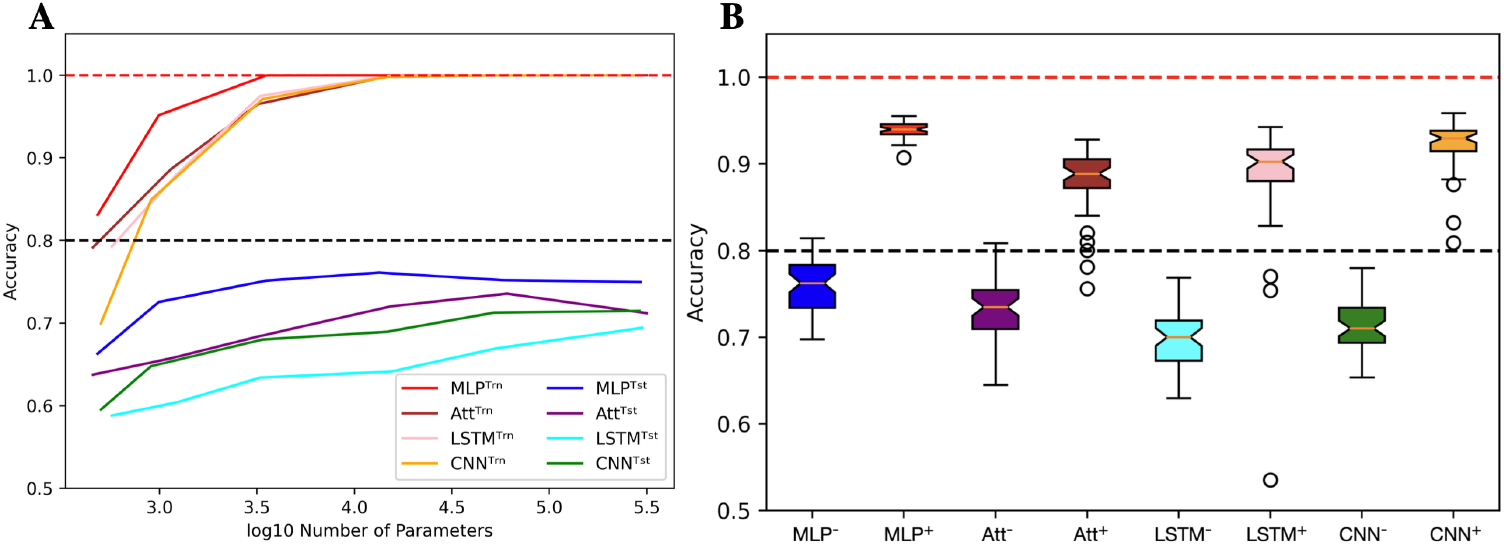
Limits when learning with few examples with mislabels in extrapolation context. (A) Influence of capacity on train (Trn) and test (tst) accuracies. The training sets are unbalanced and mislabelled as parameters (*N, L, µ, α*) are fixed to (500, 6, 0.2, 0.4). The testing sets on the opposite are balanced and correctly labelled with parameters (*N, L, µ, α*) being fixed to (4^8^, 8, 0.0, 0.5). (B) With the same parameters, distributions of test accuracies over 50 simulations are reported, with models being trained on 500 examples (e.g. MLP^-^) or 4000 examples (e.g. MLP^+^). The capacity used for each situation is the best capacity achieved on the test sets with respect to the heatmaps in Fig 8. Dotted lines indicate the 100% and 80% accuracy marks since 20% of the training examples are mislabelled.

In order to gain more insight into the behavior of the four models when dealing with few or many training examples, we compared their distributions of performances (Fig 9B). To produce this figure, we used the capacities that maximize accuracy based on the heatmaps presented in Fig 8. As a result, a wide range of capacities was needed for the models, with the MLP using *C ≈* 3500 and the CNN using *C ≈* 15 000 when trained on many examples. On the other hand, the Att and LSTM required much higher capacities, with the Att using *C ≈* 60 000 and the LSTM using *C ≈* 300 000 regardless of the number of training examples. Analyzing these distributions, we observe that the MLP model appears to be the most suitable for extrapolating with mislabels since its test accuracies have a higher mean and lesser variance than the other three models.

With such a fundamental binary classification task, the behaviors of some specific classical ML algorithms can put the results on neural networks into perspective. Focusing on the *k*-nearest neighbors (KNN), the support vector machine (SVM), the decision tree (tree) and the random forest (forest) algorithms, we measured the performances of such methods when learning on few examples with mislabels in a length-wise extrapolation context. All algorithms use default `sklearn` implementation except the KNN uses 35 neighbors, the tree uses a maximum depth of 12 decisions and the forest uses 400 classification trees. These parameters were determined by grid search maximizing the generalization performance for most training dataset sizes.

First of all, we measured the ability of these algorithms to account for mislabels in the training dataset (Fig 10A). In comparison to the four tested neural networks, the method that behaves in the most similar way is the SVM, as the test accuracy tends to 100% while the train accuracy tends to 1 *− µ* as *N → ∞*. The decision tree bahaves similarly, but its generalization performances are not as good. The KNN also performs poorly, but its train accuracies are stuck at 100%. The most surprising behavior is held by the random forest algorithm: Even tough it fits 100% of the training datasets, which include 20% of mislabelled examples, the test accuracies can simultaneously reach above 99%, which shows an ability to minimize both test and training risks [Belkin et al.(2019)Belkin, Hsu, Ma, and Mandal] [Peters and Schuld(2022)]. The accuracy distributions presented in Fig 10B allow to further visualize this diversity of behaviors when algorithms are trained with 500 and 4000 examples of length 6 before being tested on the whole set of sequences of length 8. Note that despite the relatively good performances of the SVM and random forest algorithms, they also suffer from the same misclassification problem presented in Fig 6.

**Fig 10.**
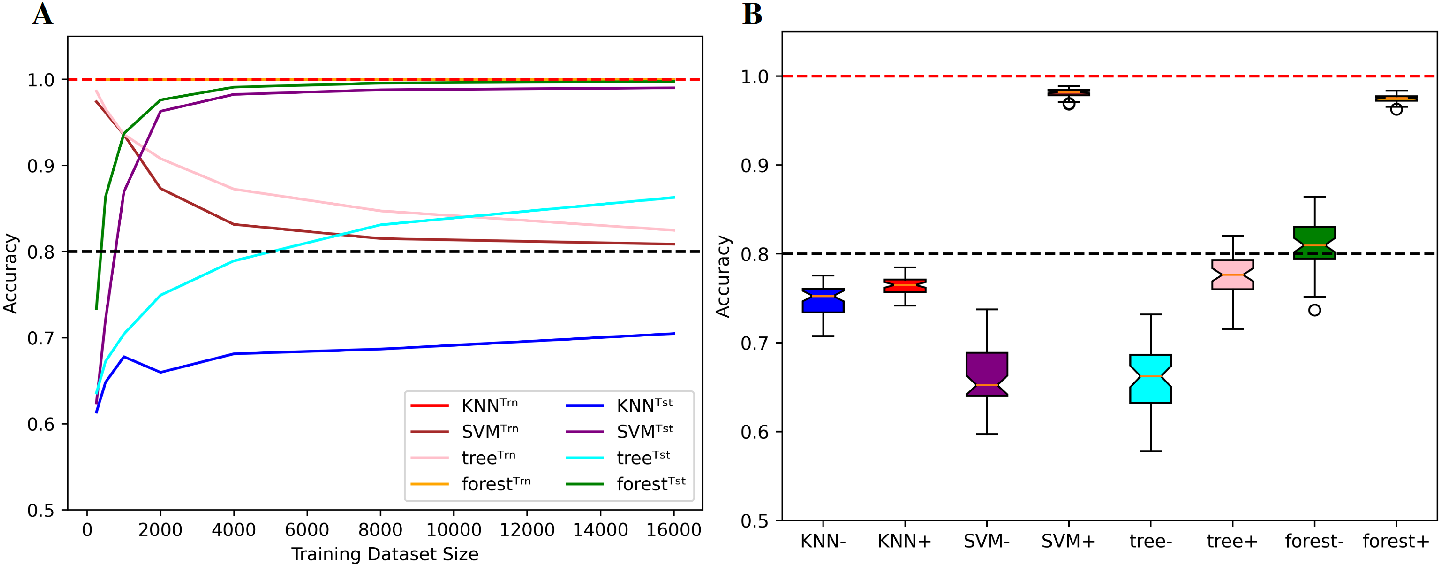
Behaviors of classical ML methods when extrapolating with mislabels. (A) Influence of the training dataset size over train (Trn) and test (Tst) accuracies for specific classical ML algorithms. Sequence length is fixed to *L* = 8 and mislabelling probability is fixed to *µ* = 0.2. (B) With (*µ, α*) being fixed to (0.2, 0.4), the models are trained on sequences of length 6 before being tested on the whole set of sequences of length 8. Distributions of test accuracies over 50 simulations are reported. The dataset size varies between *N* = 500 (e.g. KNN^-^) and *N* = 4000 (e.g. KNN^+^). Dotted lines indicate the 100% and 80% accuracy marks since 20% of the training examples are mislabelled.

Overall, this section highlights the importance of considering multiple challenges simultaneously when measuring the impact of capacity and training dataset size on the generalization performance of a variety of ML models. It also underscores the need for a better understanding of the trade-offs involved in choosing appropriate models and capacity for specific tasks, especially when dealing with small training datasets.

## Conclusion

In conclusion, the use of statistical learning through neural networks holds great promise for gaining insight into the complex mechanisms that govern RNA folding. Even though over-parameterized models may be adequate for learning useful representations, the fact remains that the quality of a ML model highly depends on the data on which it has been trained. Our study highlights three main challenges researchers face when working with current RNA structure data and provides suggestions for overcoming them.

Specifically, when dealing with mislabels, low-capacity models may be preferable, as long as enough training examples are provided. Moreover, for tasks that require extrapolating to structurally dissimilar data, high-capacity models may provide better performance. Finally, we recommend exploring the behavior of a variety of neural networks on synthetic data to better understand their specific risks and benefits in predicting RNA structure. Overall, by addressing these challenges, machine learning could provide valuable insights into RNA folding and contribute to the development of new approaches for studying biological systems involving RNA.

## Acknowledgments

We would like to express our sincere gratitude to Sébastien Lemieux and all the participants of the Computational Approaches to RNA Structure and Function conference held in Benasque, Spain in 2022 for their valuable suggestions and feedback.

## Supporting information

**S1 Fig.**
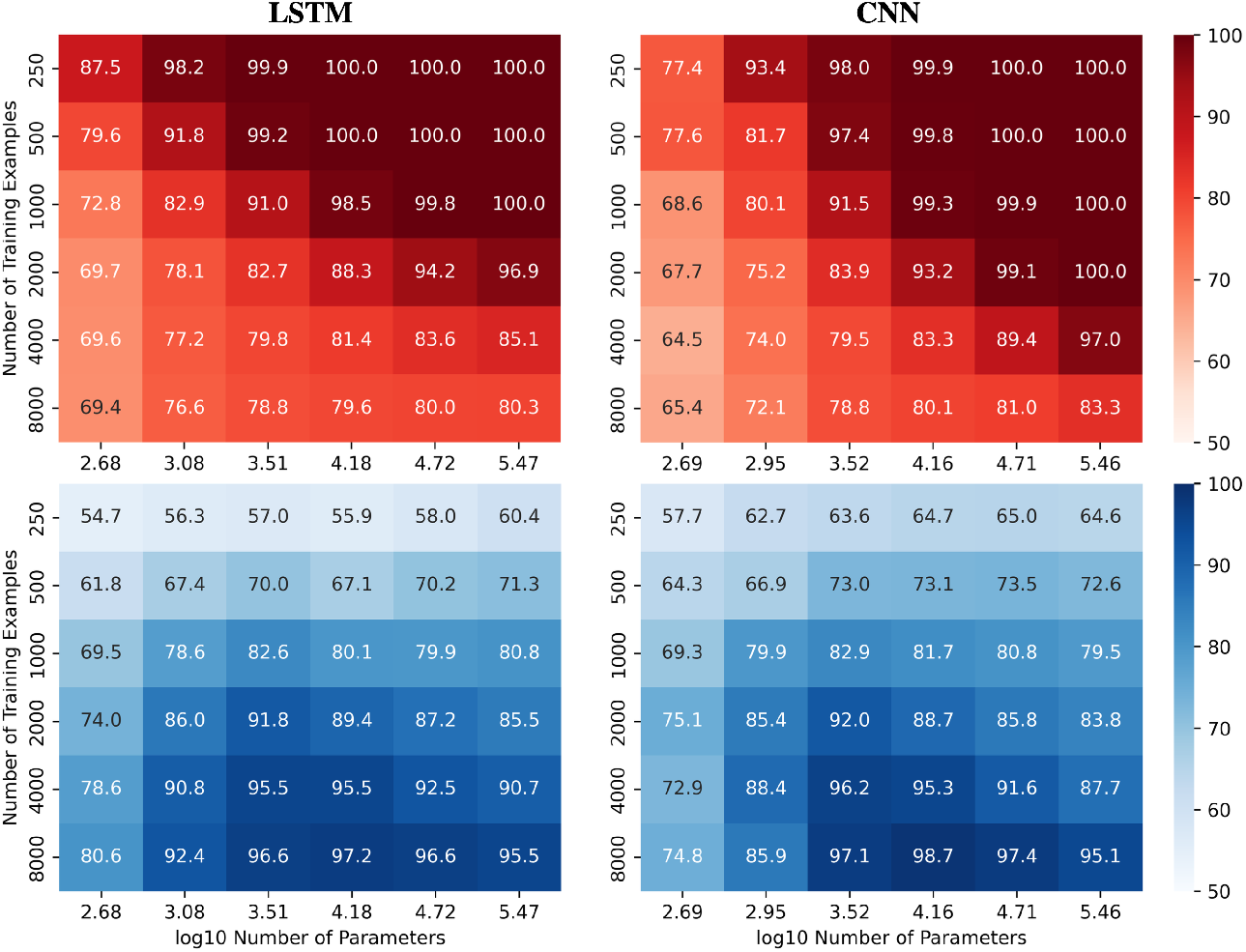
Performance of LSTM and CNN models when learning with mislabels. Train (red) and test (blue) mean accuracies over 50 simulations reported for LSTM and CNN models. Sequence length and mislabelling probability are respectively fixed to *L* = 8 and *µ* = 0.2.

**S2 Fig.**
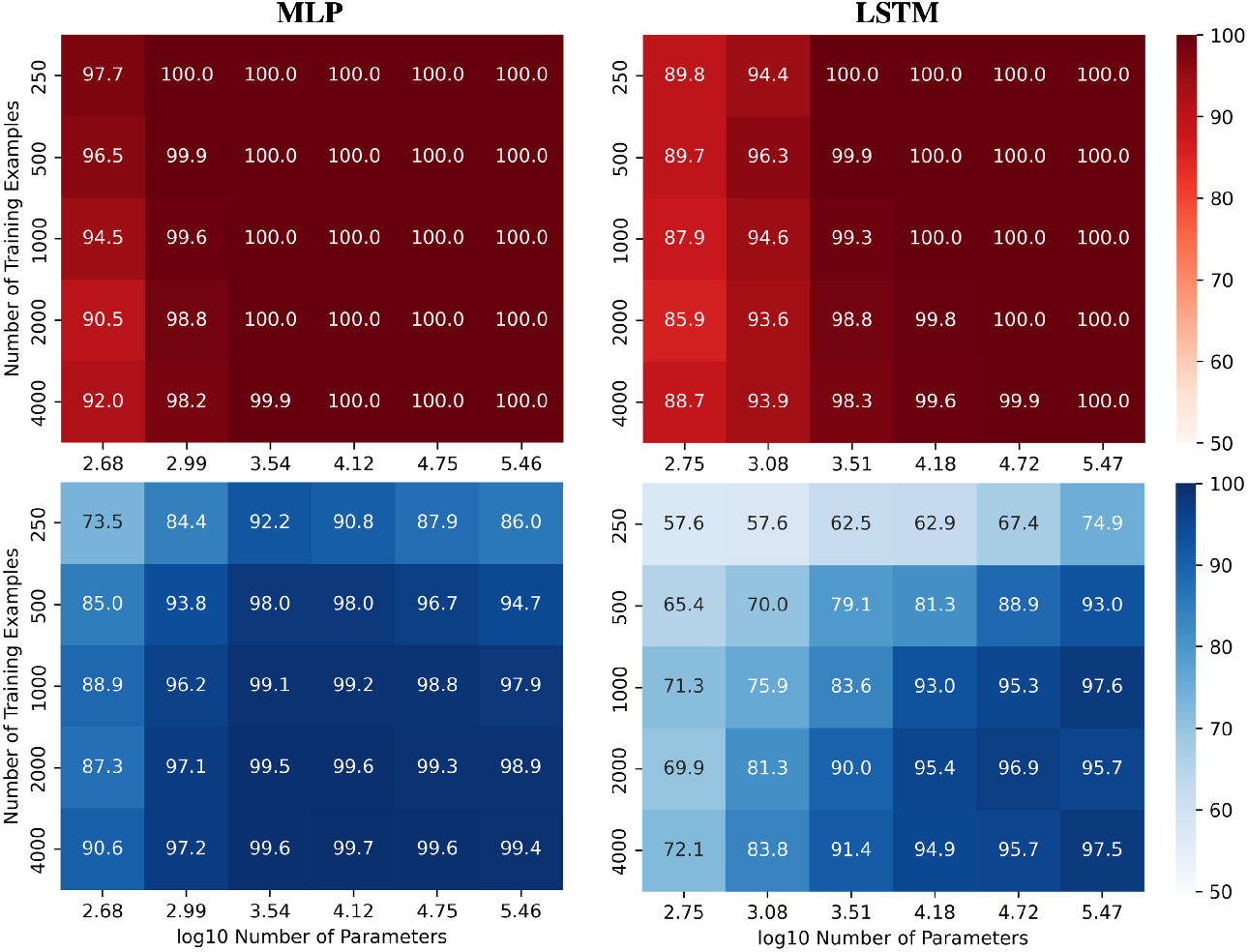
Performance of MLP and LSTM models in length-wise extrapolation context. Train (red) and test (blue) mean accuracies over 50 simulations reported for MLP and LSTM models. Models were trained on sequences of length 6 before being tested on sequences of length 8, without mislabelling in the training set.

**S3 Fig.**
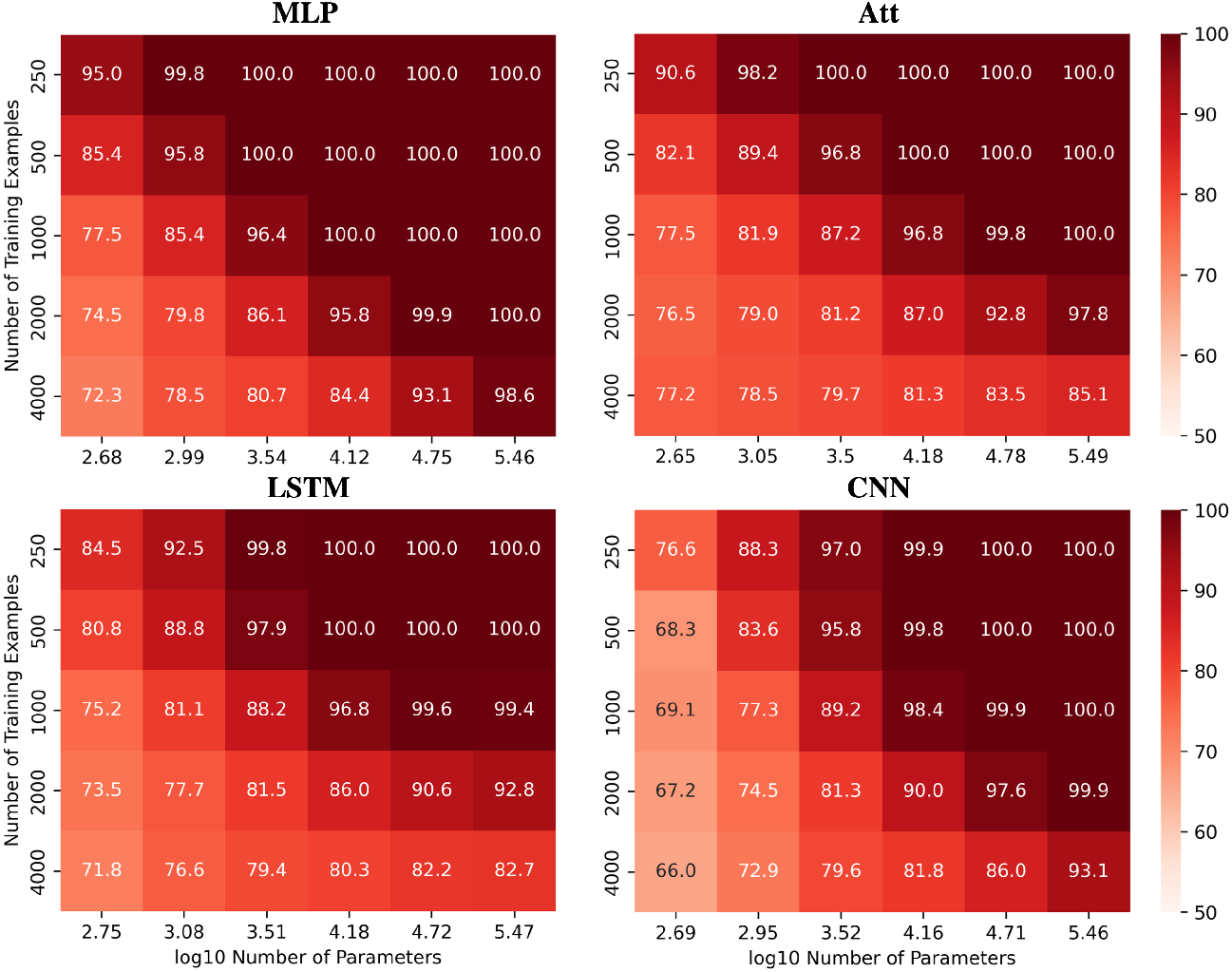
Training performances for all models when learning with mislabels in extrapolation context. Train accuracies for all models when trained on sequences of length 6 with 20% mislabelled training examples and 40% positivity rate.

## References

Balestriero, Pesenti, and LeCun Balestriero, R., Pesenti, J., and LeCun, Y. (2021). Learning in high dimension always amounts to extrapolation. arXiv preprint arXiv:2110.09485

Belkin Hsu, Ma, and Mandal Belkin, M., Hsu, D., Ma, S., and Mandal, S. (2019). Reconciling modern machine-learning practice and the classical bias–variance trade-off. Proceedings of the National Academy of Sciences 116, 15849–15854

Berrada, Zisserman, and Kumar Berrada, L., Zisserman, A., and Kumar, M. P. (2020). Training neural networks for and by interpolation. In International conference on machine learning (PMLR), 799–809

Burley, Bhikadiya, Bi, Bittrich, Chen, Crichlow et al. Burley, S. K., Bhikadiya, C., Bi, C., Bittrich, S., Chen, L., Crichlow, G. V., et al. (2022). Rcsb protein data bank: Celebrating 50 years of the pdb with new tools for understanding and visualizing biological macromolecules in 3d. Protein Science 31, 187–208

Chen, Li, Umarov, Gao, and Song Chen, X., Li, Y., Umarov, R., Gao, X., and Song, L. (2020). Rna secondary structure prediction by learning unrolled algorithms. arXiv preprint arXiv:2002.05810

Condon, Davy, Rastegari, Zhao, and Tarrant. Condon, A., Davy, B., Rastegari, B., Zhao, S., and Tarrant, F. (2004). Classifying rna pseudoknotted structures. Theoretical Computer Science 320, 35–50

Danaee, Rouches, Wiley Deng, Huang, and Hendrix Danaee, P., Rouches, M., Wiley, M., Deng, D., Huang, L., and Hendrix, D. (2018). bprna: large-scale automated annotation and analysis of rna secondary structure. Nucleic acids research 46, 5381–5394

Dumoulin, V. and Visin, F. (2016). A guide to convolution arithmetic for deep learning. arXiv preprint arXiv:1603.07285

Flamm, Wielach, Wolfinger, Badelt Lorenz, and Hofacker Flamm, C., Wielach, J., Wolfinger, M. T., Badelt, S., Lorenz, R., and Hofacker, I. L. (2021). Caveats to deep learning approaches to rna secondary structure prediction. Biorxiv, 2021–12

Fu, Cao, Wu, Peng Nie, and Xie Fu, L., Cao, Y., Wu, J., Peng, Q., Nie, Q., and Xie, X. (2022). Ufold: fast and accurate rna secondary structure prediction with deep learning. Nucleic acids research 50, e14–e14

Goodfellow, Bengio, and Courville Goodfellow, I., Bengio, Y., and Courville, A. (2016). Deep learning (MIT press)

Hinton, Srivastava, Krizhevsky, Sutskever, and Salakhutdinov Hinton, G. E., Srivastava, N., Krizhevsky, A., Sutskever, I., and Salakhutdinov, R. R. (2012). Improving neural networks by preventing co-adaptation of feature detectors. arXiv preprint arXiv:1207.0580

Hochreiter, S. and Schmidhuber, J. (1997). Long short-term memory. Neural computation 9, 1735–1780

Hofacker, Fontana, Stadler, Bonhoeffer, Tacker, Schuster et al. Hofacker, I. L., Fontana, W., Stadler, P. F., Bonhoeffer, L. S., Tacker, M., Schuster, P., et al. (1994). Fast folding and comparison of rna secondary structures. Monatshefte fur chemie 125, 167–167

Ioffe, S. and Szegedy, C. (2015). Batch normalization: Accelerating deep network training by reducing internal covariate shift. In International conference on machine learning (pmlr), 448–456

Kingma, D. P. and Ba, J. (2014). Adam: A method for stochastic optimization. arXiv preprint arXiv:1412.6980

LeCun, Y. et al. (1989). Generalization and network design strategies. Connectionism in perspective 19, 18

Marchand, Will, Berkemer, Bulteau, and Ponty Marchand, B., Will, S., Berkemer, S., Bulteau, L., and Ponty, Y. (2022). Automated design of dynamic programming schemes for RNA folding with pseudoknots. In WABI 2022 - 22nd Workshop on Algorithms in Bioinformatics (Potsdam, Germany)

Moore, P. B. (1999). Structural motifs in rna. Annual review of biochemistry 68, 287–300

Nair, V. and Hinton, G. E. (2010). Rectified linear units improve restricted boltzmann machines. In Proceedings of the 27th international conference on machine learning (ICML-10). 807–814

Peters, E. and Schuld, M. (2022). Generalization despite overfitting in quantum machine learning models. arXiv preprint arXiv:2209.05523

Rivas, E. and Eddy, S. R. (1999). A dynamic programming algorithm for rna structure prediction including pseudoknots. Journal of molecular biology 285, 2053–2068

Rivas, Lang, and Eddy Rivas, E., Lang, R., and Eddy, S. R. (2012). A range of complex probabilistic models for rna secondary structure prediction that includes the nearest-neighbor model and more. RNA 18, 193–212

Sak, Senior, and Beaufays Sak, H., Senior, A. W., and Beaufays, F. (2014). Long short-term memory recurrent neural network architectures for large scale acoustic modeling. In INTERSPEECH. 338–342

Sato, Akiyama, and Sakakibara Sato, K., Akiyama, M., and Sakakibara, Y. (2021). Rna secondary structure prediction using deep learning with thermodynamic integration. Nature communications 12, 941

Shen, Hu, Peng, Chen, Xiong, Hong et al. Shen, T., Hu, Z., Peng, Z., Chen, J., Xiong, P., Hong, L., et al. (2022). E2efold-3d: End-to-end deep learning method for accurate de novo rna 3d structure prediction. arXiv preprint arXiv:2207.01586

Singh, Hanson Paliwal, and Zhou Singh, J., Hanson, J., Paliwal, K., and Zhou, Y. (2019). Rna secondary structure prediction using an ensemble of two-dimensional deep neural networks and transfer learning. Nature communications 10, 5407

Smola, A. J. and Schölkopf, B. (2004). A tutorial on support vector regression. Statistics and computing 14, 199–222

Szikszai, Wise, Datta, Ward, and Mathews Szikszai, M., Wise, M., Datta, A., Ward, M., and Mathews, D. H. (2022). Deep learning models for rna secondary structure prediction (probably) do not generalize across families. Bioinformatics 38, 3892–3899

Vaswani, Shazeer, Parmar, Uszkoreit, Jones, Gomez et al. Vaswani, A., Shazeer, N., Parmar, N., Uszkoreit, J., Jones, L., Gomez, A. N., et al. (2017). Attention is all you need. Advances in neural information processing systems 30

Wang, Liu, Zhong, Liu, Lu, Li et al. Wang, L., Liu, Y., Zhong, X., Liu, H., Lu, C., Li, C., et al. (2019). Dmfold: A novel method to predict rna secondary structure with pseudoknots based on deep learning and improved base pair maximization principle. Frontiers in genetics 10, 143

Zakov, Goldberg Elhadad, and Ziv-Ukelson Zakov, S., Goldberg, Y., Elhadad, M., and Ziv-Ukelson, M. (2011). Rich parameterization improves rna structure prediction. Journal of Computational Biology 18, 1525–1542

Zhang, Bengio, Hardt, Recht, and Vinyals Zhang, C., Bengio, S., Hardt, M., Recht, B., and Vinyals, O. (2021). Understanding deep learning (still) requires rethinking generalization. Communications of the ACM 64, 107–115

Zhao, Zhao, Fan, Yuan Mao, and Yao Zhao, Q., Zhao, Z., Fan, X., Yuan, Z., Mao, Q., and Yao, Y. (2021). Review of machine learning methods for rna secondary structure prediction. PLoS computational biology 17, e1009291

Zuker, M. and Stiegler, P. (1981). Optimal computer folding of large rna sequences using thermodynamics and auxiliary information. Nucleic acids research 9, 133–148

